# N-formylation modifies membrane damage associated to PSMα3 interfacial fibrillation

**DOI:** 10.1101/2024.02.27.582255

**Authors:** Laura Bonnecaze, Katlyn Jumel, Anthony Vial, Lucie Khemtemourian, Cécile Feuillie, Michael Molinari, Sophie Lecomte, Marion Mathelié-Guinlet

## Abstract

The virulence of *Staphylococcus aureus*, a multi-drug resistant pathogen, notably depends on the expression of the phenol soluble modulins α3 (PSMα3) peptides, able to self-assemble into amyloid-like cross-α fibrils. Despite remarkable advances evidencing the crucial, yet insufficient, role of fibrils in PSMα3 cytotoxic activities towards host cells, the relationship between its molecular structures, assembly propensities, and modes of action remains an open intriguing problem. In this study, combining Atomic Force Microscopy (AFM) imaging and infrared spectroscopy, we first demonstrated *in vitro* that the charge provided by the N-terminal capping of PSMα3 alters its interactions with model membranes of controlled lipid composition, without compromising its fibrillation kinetics or morphology. N-formylation eventually dictates PSMα3 - membrane binding *via* electrostatic interactions with the lipid head groups. Furthermore, PSMα3 insertion within the lipid bilayer is favoured by hydrophobic interactions with the lipid acyl chains, only in the fluid-phase of membranes, and not in the gel-like ordered domains. Strikingly, our real-time AFM imaging emphasizes how intermediate protofibrillar entities, formed along PSMα3 self-assembly and promoted at the membrane interface, likely disrupt membrane integrity *via* peptide accumulation, and subsequent membrane thinning in a peptide concentration and lipid-dependent manner. Overall, our multiscale and multimodal approach sheds new light on the key roles of N-formylation and intermediate self-assembling entities, rather than mature fibrils, in dictating deleterious interactions of PSMα3 with specific membrane lipids, likely underscoring its ultimate cellular toxicity *in vivo*, and in turn *S. aureus* pathogenesis.

## Introduction

*Staphylococcus aureus* is a human commensal of the microbiota and the human epithelia, which can turn into an opportunistic pathogen, eventually causing life-threatening diseases. It is also well-known for its key role in nosocomial infections and its resistance to many antibiotics^1,2^. *S. aureus* virulence has been shown to critically depend on the production of a family of small peptides, the phenol soluble modulins (PSMs)^3,4^: the small positively-charged α-type PSMs (∼20-25 amino acids, aa) and the long negatively-charged β-type PSMs (∼44 aa). PSMs share intrinsic properties, such as amphipathy, α-helical content, and a propensity to aggregate^5–10^. All PSMs are indeed able to self-assemble, from soluble α-helical monomers to intermediate oligomeric entities of diverse sizes and shapes and insoluble unbranched fibrils. These fibrils are mostly characterized by a stacking of β-sheets where individual β-strands run perpendicular to the fibril axis and that associate *via* an extensive hydrophobic core^11,12^. Initially, this so-called cross-β scaffold was solely associated to self-assembled proteins involved in human neurodegenerative disorders (called pathological amyloids)^13–16^. But, the past two decades have witnessed the emergence of amyloids with beneficial and physiological roles (called functional amyloids)^17–19^, such as PSMs allowing *S. aureus* to circumvent *in fine* the immune defenses. Markedly, among all PSMs, only PSMα3 participates in all infection processes, from cytolytic activities, to the structuration and dissemination of biofilms, and the eventual triggering of the pro-inflammation cascade^3^. This intriguing multi-functionality might be associated to the unique and remarkable cross-α structure, where α-helices “replace” β-strands, adopted by PSMα3 fibrils^5,6^ or their cross-α/β polymorphism later revealed^20,21^.

Among other functions, PSMα3 exhibits, at micromolar concentrations, the highest toxicity of all PSMs towards eukaryotic cells^5,22–24^. Such cytotoxicity primarily depends on the dynamic interactions between PSMα3 and host cell membranes, their first targets *in vivo*. Although the exact modes of action are still unknown, PSMα3 toxicity is not mediated by specific cell receptors but putatively requires the cross-α fibrillation of the wild-type (WT) peptides: the positive charges carried by WT-PSMα3 (overall charge +2; Fig. 1A) likely mediate their interactions with cell membranes and fibrillation regulates the availability of such charges to subsequently drive cytotoxicity^22^. Based on Förster resonance energy transfer and fluorescence anisotropy experiments, Malishev *et al*. also highlighted that WT-PSMα3 fibrils could insert into eukaryotic mimetic membranes mainly composed of zwitterionic lipids, while they could only accumulate at the negatively-charged bacterial membrane interface^25^. Further complexifying a comprehensive picture of PSMα3 cytotoxicity is the co-existence *in vivo* of two forms, a formylated (f-) and deformylated (df-) form, as PSMα3 initially translates with a formyl group at the N-terminal that can be cleaved under specific conditions^26^. This formyl group not only lowers the overall charge of the peptide (+1) but modifies the N-terminal charge of the methionine, from (-C_α_ -NH-COH) upon translation to (-C_α_-NH3^+^) when cleaved. While the importance of terminal capping in amyloid fibril formation, structure and morphology is known^27,28^, and partially characterized for PSMα3^29^, the role of N-formylation in PSMs functions still remains under debate. This role has been only discussed in light of its pro-inflammation^30–32^ and biofilm scaffolding activities^33^. Besides, *in vitro* investigation on PSMα3 toxic activities have been only performed on the deformylated forms.

**Figure 1.**
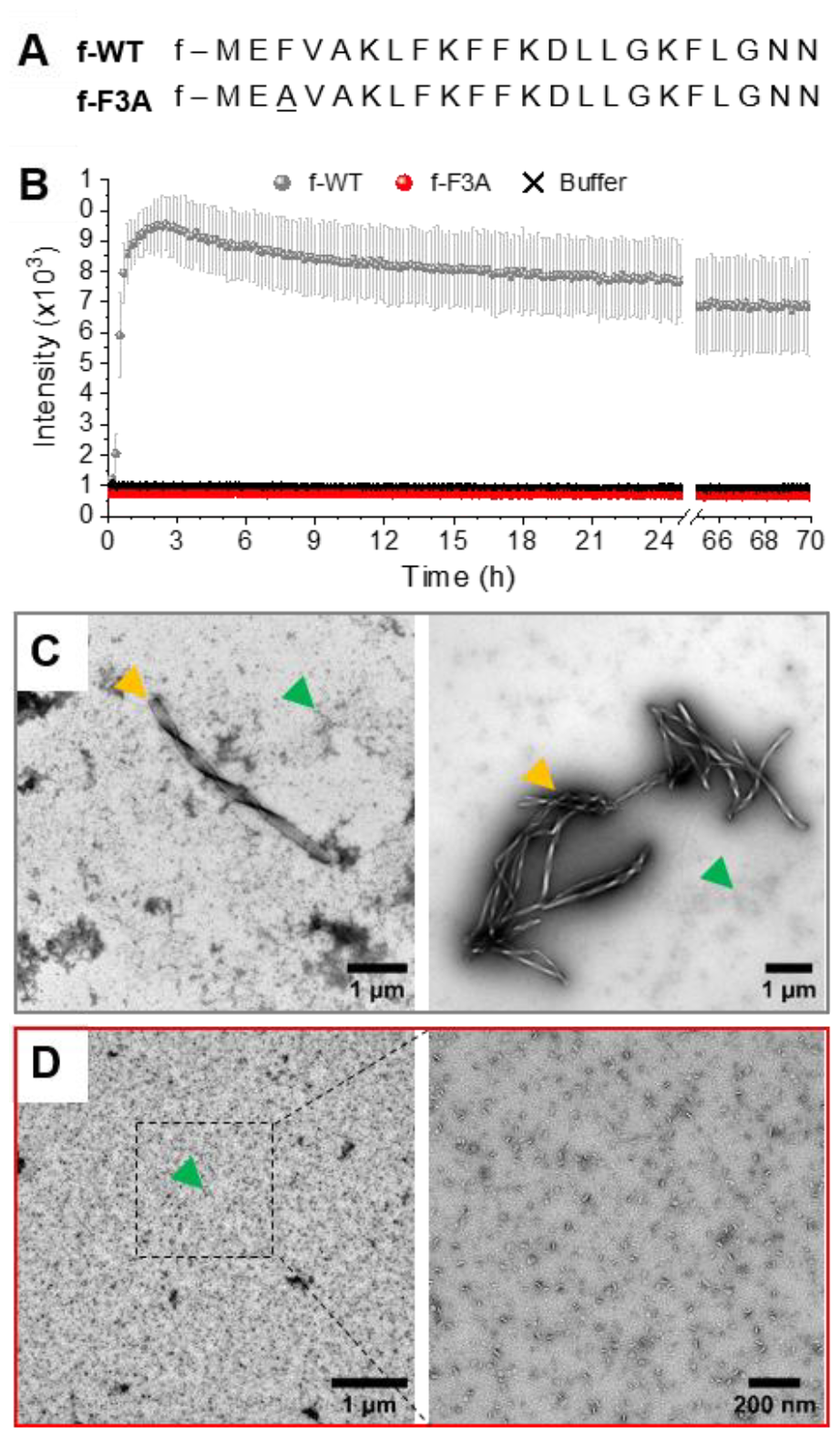
Fibrillation of formylated PSMα3. (A) Amino-acid sequences of the wild type (WT) and single-point mutation (F3A) PSMα3, with a formylation (f-) at their N-terminal. (B) Kinetics of PSMα3 fibril formation followed by ThT fluorescence, at 37°C and a peptide concentration of 50 µM. Error bars stand for the standard deviation between three replicates. (C-D) Negatively stained TEM images of PSMα3 (C, f-WT and D, f-F3A) after 3 days of incubation at 37°C. Yellow and green arrows point at mature fibrils and aggregates respectively.

The key roles of both PSMα3 intrinsic properties and cell membrane composition in the possible toxic mechanism(s) of PSMα3 have been further evidenced by the behaviour of a mutant peptide where the phenylalanine in position 3 has been replaced by an alanine (F3A, Fig. 1A). Indeed, these F3A-PSMα3 peptides, that are intrinsically unable to form amyloid fibrils, induce a much reduced toxicity towards human cells^5,22^, supporting the importance of cross-α fibrillation in the deleterious activity of PSMα3. Unexpectedly, they also lead to bactericidal activities towards some gram-positive bacteria^6,34–36^, a gained function compared to WT-PSMα3, highlighting the species-specific toxicity of PSMα3. An *in vitro* study additionally showed that df-F3A-PSMα3 could actually aggregate on bacterial mimetic membranes, only as small oligomers deposited at the interface^25^. Consequently, the reduced cytotoxicity of F3A-PSMα3 and its antibacterial activity could not be correlated to any fibril insertion within the cell membranes, suggesting that monomers and/or oligomers, independently of amyloid fibrillation, could also trigger membrane destabilization, in a lipid-dependent manner. Interestingly, for pathological amyloids (*e*.*g*. Aβ), the amyloid cascade hypothesis, according to which insoluble fibrils cause cell toxicity^37^, is increasingly challenged as intermediate oligomeric entities, formed during the self-assembly process, were reported to drive cell membrane disruption, and subsequent cell dysfunction and pathogenesis^38–40^.

Over the last decade, an increasing knowledge about the structure and cytotoxic activities of PSMα3 has been gained, especially since the discovery of its unique cross-α fibril structure. However, due to the lack of integrative highly resolved techniques, amyloid(-like) fibrillation and toxicity were only considered independently, without direct correlation between the morphological and structural states of PSMα3 and their biological functions *in fine*. Thus, because of the high dynamics at play both at the cell membrane interface and in between entities co-existing in solution (exchange between mono-, oligomers and fibrils), it still remains elusive which, and how, entities formed along PSMα3 self-assembly mediate interactions with cells. Moreover, the role of N-formylation in the possible deleterious activities of PSMα3, and the underlying molecular mechanisms, have not been addressed so far. Here, we present an *in vitro* molecular investigation of the interactions between PSMα3, in their formylated form (both f-WT and f-F3A), and supported lipid bilayers (SLB) of controlled lipid composition. These biomimetic systems allow (i) to overcome the high complexity of living cells, in physiologically-relevant conditions^41^ and (ii) to disentangle how the physico-chemical properties of both interacting partners lead to specific modes of action of PSMα3. Combining highly-resolved imaging and spectroscopic techniques, namely atomic force microscopy (AFM) and Attenuated Total Reflectance Fourier Transform infrared spectroscopy (ATR-FTIR), morphological, mechanical, and structural features of the peptide-membrane interactions have been revealed from the micro- to the nanoscale, in real-time. Our results first indicate that N-formylation impacts the self-assembly propensity of PSMα3 at the membrane interface, in a lipid-dependent manner. Strikingly, they further demonstrate that oligomeric entities are likely the membrane-active entities that tend to insert only in the fluid phase of complex membranes, eventually disrupting their integrity and underscoring the species-specific toxicity of PSMα3. Such mechanistic insights into the molecular mechanisms developed by the secreted PSMα3 toxin are key in the efficient drug development strategies that currently focus on targeting virulence determinants of *S. aureus* to elicit less resistance than traditional antibiotics.

## Results

### Fibrillation of formylated PSMα3

The self-aggregation of f-PSMα3 was first followed by ThT fluorescence, a widely used technique to monitor amyloid fibrillation *via* the increase in fluorescence, ThT only binding to hydrophobic cavities within amyloid(-like) structures^42,43^. As illustrated by the typical sigmoidal curve, f-WT self-assembles into amyloid structures at 37°C, and at a concentration of 50 µM in 10 mM sodium phosphate buffer complemented with 150mM NaCl (pH 8.0) (Fig. 1B). This self-assembly rapidly occurs with a very short lag phase (<1h) followed by a sharp transition to the final plateau reached within two hours. This kinetics does not critically depend on the peptide concentration (Fig. S1). Amyloid structures formed by f-WT after 3 days at 37°C were characterized by Transmission Electron Microscopy (TEM) as thick fibrils, often twisted and clustered into bundles, and co-existing with more globular aggregates (Fig. 1C). These results are consistent with the intrinsic propensity of PSMα3, either formylated or deformylated, to form amyloid fibrils as previously reported^5,29^. We additionally performed such characterization for the single-point mutant F3A. f-F3A, showed no ThT binding (Fig. 1B), and remained as small aggregates even after 3 days at 37°C (Fig. 1D), suggesting that f-F3A, like df-F3A, cannot assemble into amyloid structures^5^. Together, these results demonstrate that the N-terminal formylation does not change the intrinsic capacity of WT-PSMα3 to fibrillate or the incapacity of F3A-PSMα3 to self-assemble. For WT-PSMα3, formylation has also no impact on the kinetics of fibrillation or the morphology of the formed fibrils (see Fig. S2 for a comparison between f- and df-WT). Of note, df-WT displays a higher ThT fluorescence intensity than f-WT, suggesting that both forms could differ either in their intrinsic amyloid structures or quantity of formed fibrils.

### Formylated PSMα3 accumulates on and eventually disrupts membranes

In light of its potential cytotoxic activities previously reported in the literature, we then wondered how f-WT behaves in the presence of the first cellular barriers, namely the cellular membranes. To reflect those peptide-membrane interactions, we investigated *in vitro* the structural and potential deleterious effect of f-WT when interacting with SLB of controlled composition. While zwitterionic lipids are used in combination with sphingomyelin (SM) and cholesterol (Chol) to mimic eukaryotic membranes (DOPC/SM/Chol (67:8:25)^44^), negatively charged lipids are used to reflect bacterial membranes (DOPE/DOPG (1:1)^45^). Peptides were incubated 3 days at 37°C to reach amyloid fibrillation, as determined above, and the final solution was injected on those SLBs.

Polarized ATR-FTIR first allowed to assess both the organization of the lipid membrane and the structural evolution of the peptides^46,47^ following a 3h incubation of 10 µM f-WT at the membrane interface. The membrane organization can be determined *via* the analysis of the antisymmetric and symmetric stretching modes of CH_2_ of the lipid tails (v_s_ (CH_2_) ∼ 2855 cm^-1^ and v_as_ (CH_2_) ∼ 2925 cm^-1^) (Fig. 2A). While the wavenumbers of these modes notably illustrate the fluidic properties of the membrane, the absolute absorbance at a specific wavenumber (*e*.*g*. v_as_) reflects the amount of lipids attached to the sensor and allows to validate the formation of a single lipid bilayer. Importantly, to rule out the eventuality of membrane intrinsic instabilities over the duration of peptide treatment, spectra of pure SLB have been recorded over 3 h, and the lipid coverage reproducibly determined after 3 h (Fig. 2B) is thereafter compared to the one following peptide addition. Comparisons of spectra acquired before and after f-WT addition revealed that f-WT could induce membrane damage, lipid depletion events being evaluated as a decrease in the lipid coverage of the sensor (Fig. 2A). While f-WT did not alter the lipid coverage of eukaryotic mimetic membranes (DOPC/SM/Chol), it induced a slight depletion of pure DOPC membranes (from 93 ± 1 to 77 ± 9 % coverage) (Fig. 2B). This could suggest that the peptides mainly impact the fluid phase of the membrane or, alternatively, that the more rigid / ordered phase (here enriched in cholesterol) could protect the membrane from f-WT deleterious activity.

**Figure 2.**
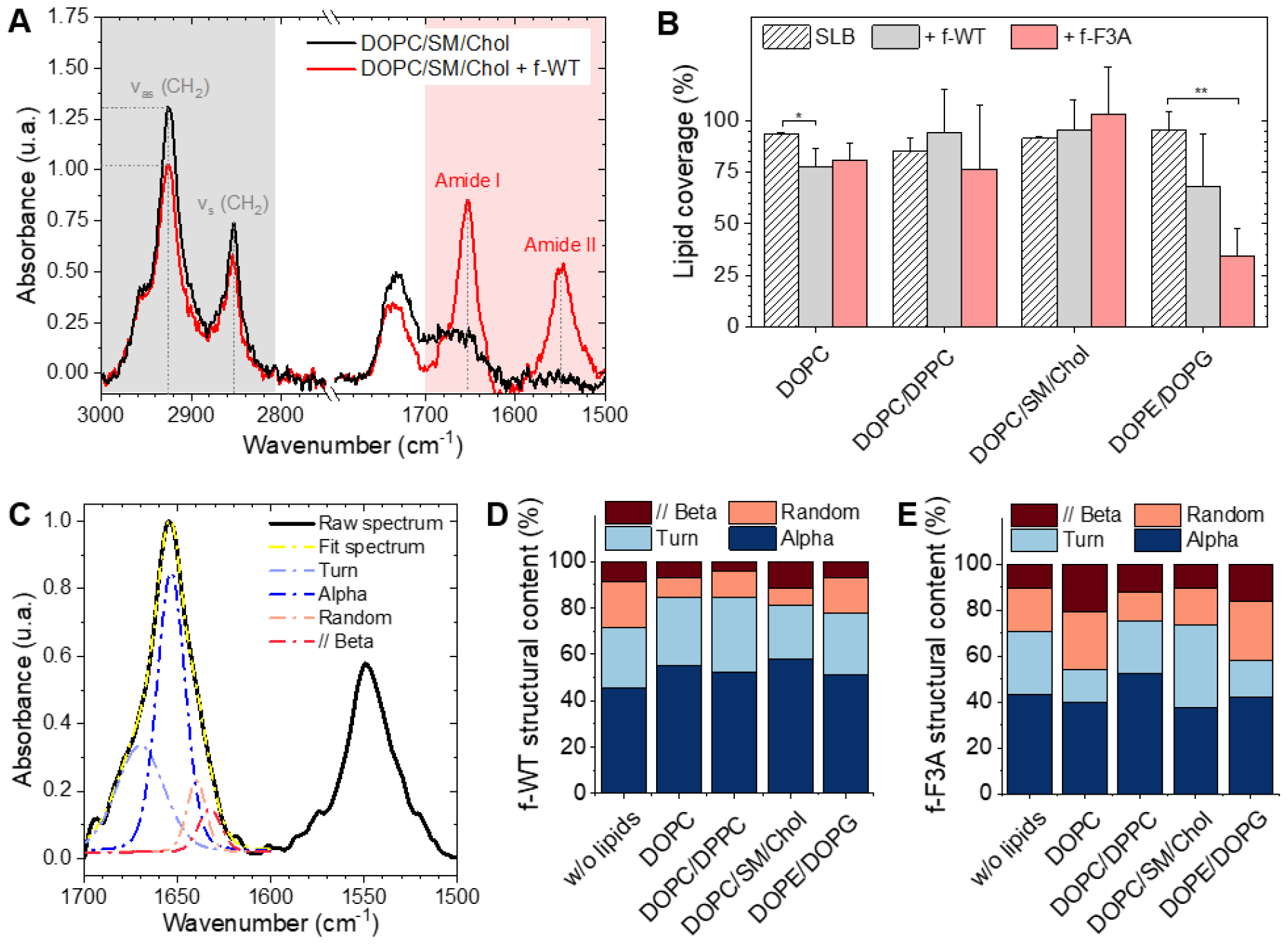
Membrane disruption via f-PSMα3 accumulation. (A) ATR-FTIR spectra of a DOPC/SM/Chol SLB before and after a 3 h incubation with f-WT at 10 µM, focused on the CH2 lipid and Amide bands. (B) Lipid coverage of the ATR-FTIR sensor, determined via the variations in vas (CH2) intensity, for SLB of diverse composition following PSMα3 addition (3 h, 10 µM). Results are presented as mean ± standard deviation of at least three independent replicates. * p < 0.05, ** p < 0.01, *** p < 0.001. (C) Deconvolution of the ATR-FTIR spectra for the Amide range of a DOPC SLB after a 3 h incubation with f-WT at 10 µM. (D-E) Secondary structure content derived from the ATR-FTIR spectra of (D) f-WT and (E) f-F3A after interacting with different SLB for 3 h at 10 µM. All spectra presented, and the resulting analysis, were obtained in the p-polarization.

To validate this hypothesis, we performed an additional control with a binary mixture of DOPC and DPPC, DPPC providing a rigid and more ordered scaffold for the membrane, as revealed by the lower wavenumber of the v_as_ (CH_2_) mode with the increasing concentration of DPPC (Table 1, Fig. S3). Similarly to eukaryotic membranes, lipid depletion was not observed after f-WT addition (Fig. 2B), even when varying the concentration of DPPC from 25 % to 75 % (Fig. 2B, Fig. S3). This reinforces the idea of a membrane fluidity-dependent action of f-WT peptides. Concerning the bacterial mimetic models (DOPE/DOPG), damage of the membrane was observed following f-WT incubation (decrease from 95 ± 9 to 68 ± 25 % in lipid coverage). Given the known cytotoxic activity of PSMα3, at micromolar concentrations and in their deformylated forms, one would expect similar functions for the formylated form herein studied and, thus, stronger impact of f-WT on SLB reflecting the composition of cellular membranes encountered *in vivo*. Such discrepancies could be explained by a dose-and time-dependent effect of PSMα3, both parameters differing when investigating *in vivo* and *in vitro* processes. Indeed, and consistent with toxicity assays on living cells and *in vitro* leakage of vesicles containing zwitterionic lipids (with varying concentration of cholesterol)^48,49^, we found that an increased concentration of f-WT peptides (50 µM) caused a drastic disruption of DOPC-containing membranes after a 1h-incubation (Fig. S4). Unexpectedly though, at this high concentration, f-WT also strongly disrupts bacterial membranes, in disagreement with the absence of antibacterial activity of the deformylated form of PSMα3.

**Table 1.** PSMα3 barely compromise membrane organization and fluidity. Wavenumber and dichroic ratio of the CH_2_ groups of different SLB, before (-) and after (+) peptide incubation.

|  | Peptides | f-WT |  | f-F3A |  |
| --- | --- | --- | --- | --- | --- |
| | | $\nu_{as}$ (CH<sub>2</sub>) (cm<sup>-1</sup>) | $R_{ATR}(\nu_{as} \text{ (CH}_2\text{)})$ | $\nu_{as}$ (CH<sub>2</sub>) (cm<sup>-1</sup>) | $R_{ATR}(\nu_{as} \text{ (CH}_2\text{)})$ |
| DOPC | - | 2925 | $1.26 \pm 0.06$ | 2924 | $1.21 \pm 0.07$ |
| | + | 2924 | $1.28 \pm 0.16$ | 2925 | $1.25 \pm 0.31$ |
| DOPC/DPPC (1:1) | - | 2920 | $1.20 \pm 0.08$ | 2921 | $1.22 \pm 0.09$ |
| | + | 2921 | $1.36 \pm 0.15$ | 2920 | $1.33 \pm 0.26$ |
| DOPC/SM/Chol (67:8:25) | - | 2925 | $1.27 \pm 0.07$ | 2924 | $1.22 \pm 0.10$ |
| | + | 2925 | $1.28 \pm 0.04$ | 2925 | $1.23 \pm 0.09$ |
| DOPE/DOPG (1:1) | - | 2924 | $1.23 \pm 0.06$ | 2925 | $1.34 \pm 0.14$ |
| | + | 2924 | $1.11 \pm 0.17$ | 2925 | $1.40 \pm 0.48$ |

To examine the role of fibrillation in such impacts on membranes, we performed the same experiments with the mutant f-F3A, also incubated for 3 days at 37°C. Similar observations were made with those peptides injected at 10 µM. They hardly affected the DOPC-containing membranes but induced a critical depletion in the bacterial ones (from 95 ± 9 to 35 ± 13 % coverage) (Fig. 2B). At higher concentration (50 µM), despite a decrease in SLB coverage, f-F3A did not significantly disrupt eukaryotic membranes or the biphasic DOPC/DPPC models, thus opposed to f-WT behaviour and consistent with the much-reduced cytotoxic activity of F3A compared to WT (Fig. S4). Besides, f-F3A induced a strong perturbation of bacterial membranes both at low (10 µM) and high concentration (50 µM), in agreement with its antibacterial activity. Noteworthy, when the membranes were still significantly constituted (lipid coverage > 25%), thus mostly after a peptide treatment at 10 µM, and whatever their lipid composition, both f-WT and f-F3A did not overall change the configuration, nor the mobility and packing of the acyl chains, as revealed by identical wavenumbers of v_as_ (CH_2_) before and after peptide addition (Table 1). Despite some slight, yet not significant, changes in the dichroic ratio of v_as_ (CH_2_) (Table 1), both peptides thus seem to preserve the initial organization and fluidity of the membranes, provided their deleterious impact is not drastic as observed at an elevated concentration of 50 µM.

Interestingly, along with the above-mentioned effects of peptides on the membrane structure, ATR-FTIR revealed that both f-WT and f-F3A specifically bind and accumulate at the membrane interface, as shown by the presence of Amide I and Amide II bands after a 3h-incubation at 10 µM (Fig. 2A). Of note, while for f-WT, the Amide I and Amide II bands were observed in most of the spectra, these bands were more rarely observed for the mutant f-F3A, yet with similar absolute absorbance as observed for f-WT (Fig. S5). Though not quantified *via* ATR-FTIR, such an observation might suggest a higher avidity of f-WT for the membranes, whatever their lipid composition, as compared to its mutant f-F3A but similar affinities of both peptides for the lipids. When the amide I band (1600-1700 cm^-1^) is significantly present, its analysis allowed to assess the secondary structure element content of the peptides bound to the membrane (Fig.2C-E). Comparing spectra obtained in a lipid-free environment and those acquired at the SLB interface revealed that the Amide I band shape did change, thus highlighting rearrangements of the peptide structure when interacting with lipids (Fig. 2D-E). Consistent with the literature on WT-PSMα3, monomeric f-WT intrinsically adopts a α-helical conformation (45 %), with additional significant contributions of turns (27 %) and random coil structures (15 %), as well as a minor proportion of parallel β-sheets (13 %) (Fig. 2D). When bound to DOPC-containing membranes, the α-helical content tends to be significantly promoted (above 53 %) and the proportion of random coil significantly reduced (below 11 %), pointing to a structuration of the peptides at the lipid interface (Fig. 2D). For instance, after interacting with eukaryotic membranes (DOPC/SM/Chol), f-WT displays 58 % of α-helical content, 23 % of turns, 7 % of random coil structures, and 12 % parallel β-sheets. This structuration into α-helical structures is less pronounced when interacting with negatively charged DOPE/DOPG membranes: f-WT exhibits 51 % of α-helical content, 26 % of turns, 16 % of random coil structures and 7 % parallel β-sheets. These results might suggest a conformational reorganization of the f-WT peptides dependending on the charge surface of the membranes. As the mutant f-F3A is concerned, it mainly adopts an α-helical conformation (43 %), with contributions of turns, random coil structures, and parallel β-sheets with similar probabilities as f-WT (Fig. 2E). Structuration of f-F3A at the membrane interface, in a lipid-dependent manner, is not as significant as for f-WT: while DOPC/DPPC and DOPC/SM/Chol membranes seem to slightly favour α-helical and turn structures, the structural content of f-F3A on other SLBs seems overall unchanged.

### Formylated PSMα3 fibrillation is favoured by DOPC

To understand the peptide-membrane interactions, and go beyond the average information obtained by polarized ATR-FTIR on the SLB at the global scale, we performed AFM high-resolution imaging experiments. Scanning SLB in real-time after peptide injection allowed to simultaneously probe potential peptide aggregation and membrane reorganization, at the nanoscale, in physiological conditions (*e*.*g*.^50,51^). To that end, provided the SLB was stable, *i*.*e*. its morphology did not change over time, the peptides were injected at a final concentration of 5 µM, slightly reduced compared to ATR-FTIR experiments for technical reasons.

The ternary DOPC/SM/Chol mixture formed a uniform and homogeneous bilayer on the mica surface, with a consistent thickness of 4-5 nm, as revealed by the presence of defects in the SLB (Fig. 3A). Following f-WT addition, the membrane morphology remains unchanged for 30-45 minutes, a period that can slightly change depending on the experiments. After this lag time, deposition and accumulation of peptides at the membrane interface are observed, as well as in the mica defects (if existing), mainly as globular aggregates or short fibrils from 1 to 5 nm thick (Fig. 3A, yellow arrows) With time, these structures grow and elongate as thin and long fibrils from 5 to 10 nm thick (Fig. 3A, blue and green arrows) until the scanned region is mostly / fully covered. This 3 steps process was reproducibly observed on different samples and does not result from tip scanning artefacts, as different areas featured the same final morphology, even if they were not scanned in real time. Similar observations were made on pure DOPC membranes. However, the phenomenon never occurred on DOPE/DOPG membranes: similar morphology (topography and roughness) of the SLB was observed in time, even after a 3h incubation with f-WT. These data first suggest that, at the local scale, f-WT interacts preferentially with DOPC-containing membranes, notably eukaryotic mimetic ones, rather than negatively charged bacterial membranes on which peptide deposition has never been observed (Fig. 3B). Markedly, the initial deposition on the SLB only featured small aggregates and/or short fibrils, able to subsequently elongate, and substantially differing from the thick clustered fibrils observed by TEM (Fig. 1B). This observation unambiguously reveals the co-existence of both mature and thick fibrils and oligomeric, if not monomeric, entities within the f-WT solution, despite the 3 days incubation at 37°C which allowed to reach saturation of amyloid formation (as shown by ThT fluorescence, Fig. 1). Moreover, these oligomeric entities appeared as nucleation spots on the SLB for the propagation of thin and elongated fibrils, suggesting the ability of zwitterionic DOPC to promote amyloid fibrillation, at solid interfaces (Fig. 3B). Interestingly, controls on DOPC/DPPC bilayers highlighted the presence of fibrils only in the liquid phase of DOPC (Fig. 3A). The gel-phase domains of DPPC, ∼ 0.5 - 1 nm thicker than the surrounding DOPC phase, remained intact upon interactions with f-WT. The real-time imaging showed that f-WT deposition and aggregation actually occurs within – and is restricted to – the DOPC phase, the growing fibrils even adopting the shape of the DPPC domain edges.

**Figure 3.**
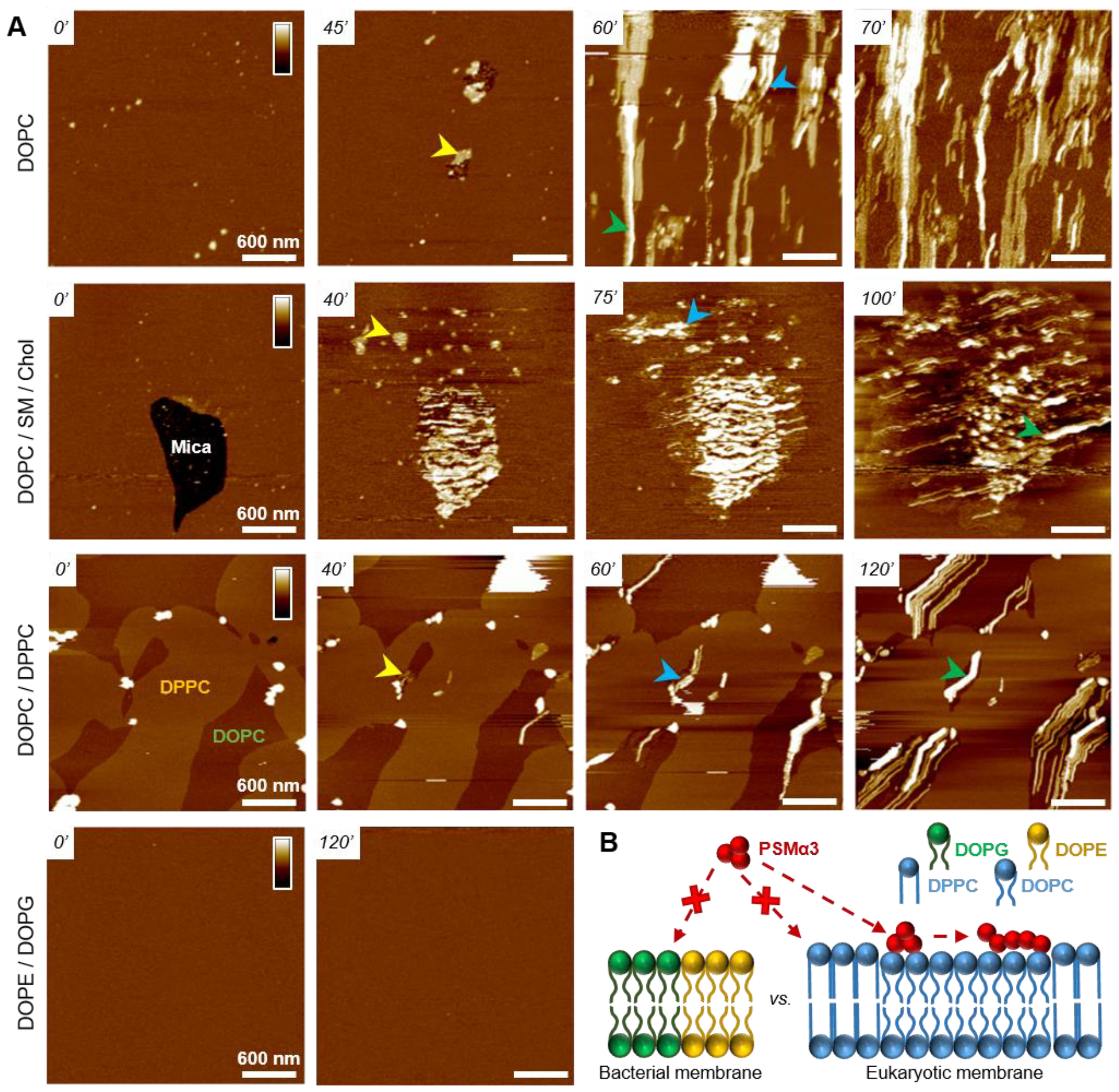
Fibrillation of f-WT PSMα3 on DOPC-containing membranes. **(A)** AFM topography images, at representative timepoints, of various SLB interacting with 5 µM f-WT. Small aggregates (yellow arrows) first appear before elongating as thin (blue arrows), and sometimes stacked (green arrows), fibrils in the fluid DOPC phases. Color scale bar: 20 nm for DOPC-SLB, 10 nm for the other SLBs. **(B)** Schematic representation of the lipid-dependent affinity, and possible fibrillation, of f-WT at the membrane interface.

These nanoscale observations confirm the affinity of f-WT for DOPC-containing SLB and the role of membrane fluidity in eventually dictating peptide-membrane interactions (Fig. 3B), consistently with our ATR-FTIR data. They additionally highlight that (i) the compact and ordered organization of DPPC prevents the f-WT accumulation, and (ii) the hydrophobic edges between gel-like DPPC domains and fluid DOPC phase potentially drive fibrils elongation and shape. They finally provide a direct visualization of the timely evolution of peptide morphology, not accessible by other techniques at this resolution: f-WT peptides evolve from small aggregates initially bound to the SLB to long and thin fibrils covering the membrane. This morphological transition reinforces the structural rearrangements determined *via* ATR-FTIR.

### Oligomeric entities of PSMα3 eventually insert and disrupt membranes

While those AFM observations emphasized the lipid-dependent fibrillation of f-WT, they also highlighted a reciprocal impact of the peptide on the SLB. Mostly on the eukaryotic and biphasic DOPC/DPPC bilayers, local damages were regularly observed as a membrane thinning, if not lipid depletion (Fig. 4).

**Figure 4.**
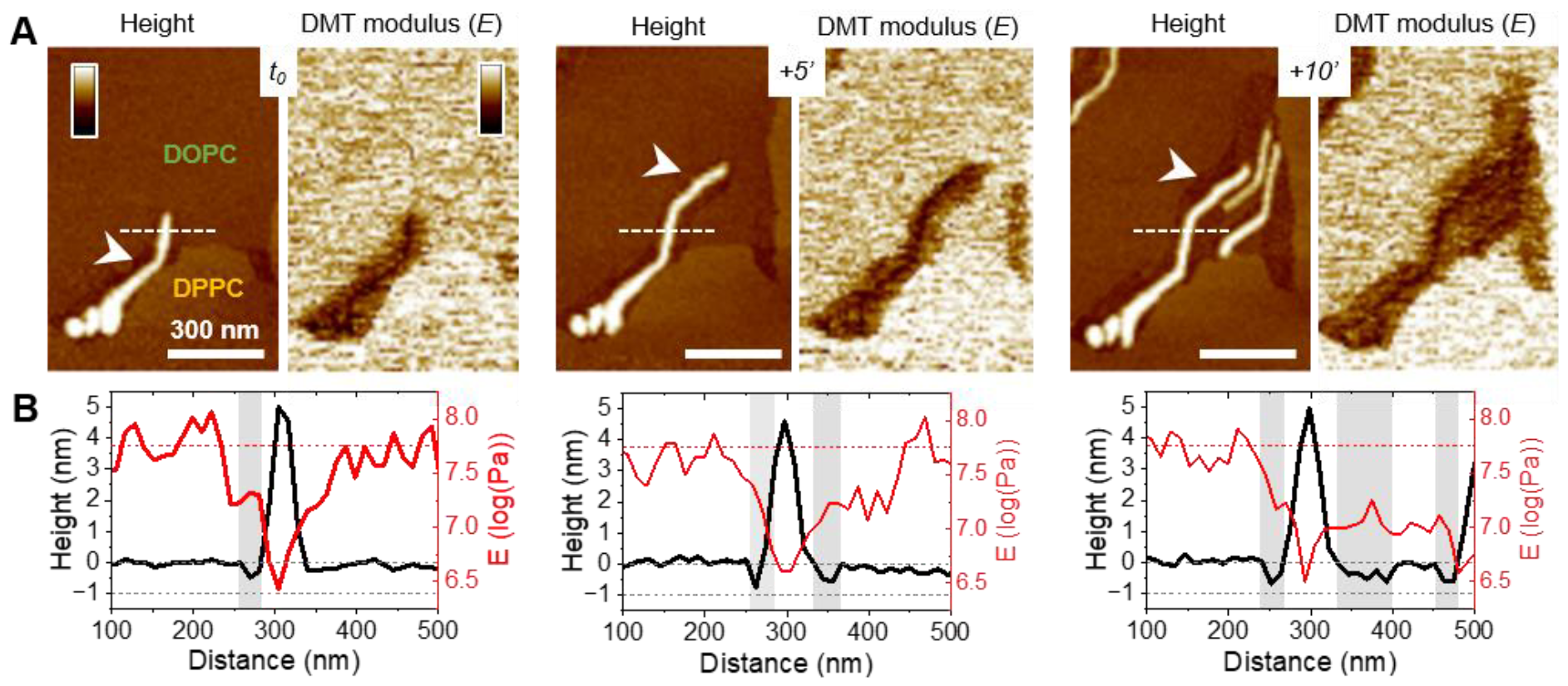
Membrane disruption by f-WT PSMα3 fibrils. Topographical and mechanical analysis of the DOPC/DPPC membrane, as an f-WT fibril elongate in time. **(A)** AFM topography and DMT modulus images, and **(B)** the corresponding height and modulus profiles, along the dashed lines, point to a membrane thinning (white arrows and grey areas) around the growing soft fibril. AFM images were obtained after f-WT, injected at 5 µM, starts interacting with – and accumulating – at the SLB interface. Color scale bars: 10 nm for the topography images and 2 log (Pa) for the DMT modulus images.

Upon peptide binding and fibrillation, an area thinner than the fluid phase (∼ 0.5 - 1 nm) was observed around the elongating fibril, and both this area and the fibril are concomitantly growing (Fig. 4A, Fig. S6). This specific area was also revealed by the nanomechanical imaging evidencing different DMT modulus: it appears softer than the lipid membrane, and barely stiffer than the fibril itself (Fig. 4A). While in some cases, those areas were smooth and homogeneous, suggesting a membrane thinning effect or a partial lipid depletion induced by the peptide (Fig. 4A), in other samples they seemed to be filled with peptidic material (Fig. 5). Indeed, part of the area could either appear “fuzzy” with small aggregates or covered with thin filaments protruding from 0.5 to 2 nm above the SLB (Fig. 5A-B, yellow arrows), thus much thinner than the mature fibrils (∼ 5 nm in height) also observed in the same areas (Fig. 5A-B, blue arrows). As for mature fibrils, these filaments, thereafter named protofibrils, grow in time and altogether form a soft area (Fig. 5C). In this area, in between fibrils and/or protofibrils, a decrease in height of approximately 0.5 - 1 nm was observed compared to the surrounding lipid membrane, and less frequently holes of ∼ 2 nm deep (Fig. 5A). When the area is densely populated with protofibrils, resulting in a typical wavy pattern, those variations were less pronounced (Fig. 5B-C). These variations in the SLB thickness might be due to the insertion of the f-WT peptides within the membrane, as also supported by the ATR-FTIR data, locally disturbing its organization and eventually leading to the disruption of the outer monolayer of the SLB. Such membrane damage following f-WT action is further supported by the disruption of SLB incubated with a more concentrated f-WT solution (> 15 µM) (Fig. S7). In the same way as at low concentration, short fibrils first deposit, and subsequently grow in/on the liquid phase of the SLB, locally disrupting the membrane as revealed by a 2 nm thinning. Then, these thinner areas progress and change the shape of the 0.5-1nm thicker DPPC domains, until they totally disappear, suggesting that both liquid and gel-like phases of the membrane are depleted by f-WT at elevated concentration, which is consistent with the significant decrease in lipid coverage measured by ATR-FTIR.

**Figure 5.**
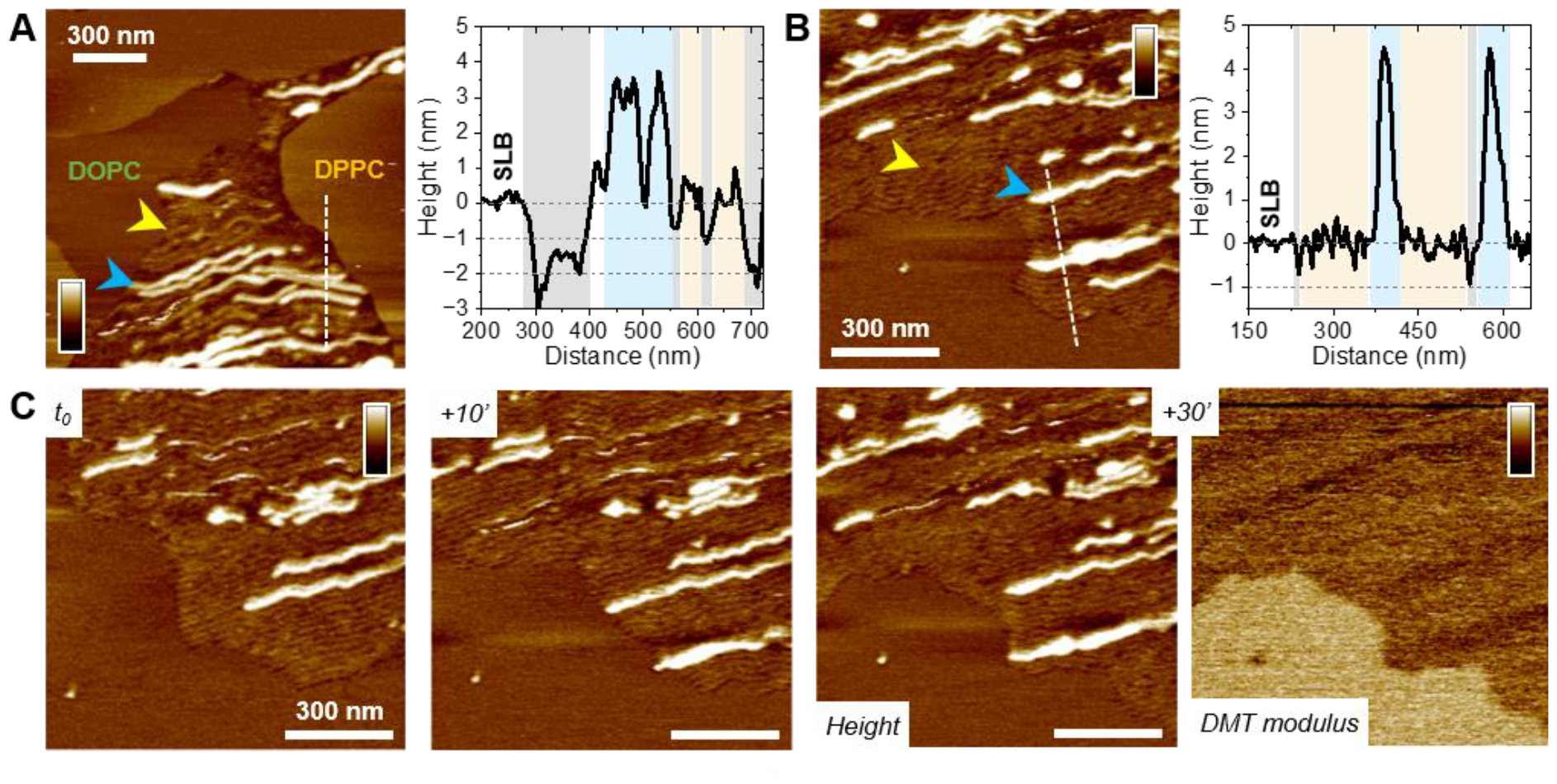
Insertion of f-WT PSMα3 protofibrils in the membrane. Topographical analysis of **(A)** DOPC/DPPC and **(B)** DOPC/SM/Chol membranes, at a representative timepoint when f-WT (injected at 5 µM) co-exist as protofibrils (yellow arrows) and mature fibrils (blue arrows). Height profiles along the dashed lines in the AFM topography images show both entities (corresponding colored areas), as well as holes and/or membrane thinning (grey areas). (C) Elongation of f-WT protofibrils, as a thin and soft « front », probed by AFM topography and DMT modulus images. Color scale bar: (A) 10 nm, (B-C) 6 nm and 2 log(Pa).

Importantly, the presence of small aggregates and protofibrils, still able to grow and elongate in time at the interface of - and even within - the liquid phase of the DOPCmembranes, suggests the co-existence of oligomers and mature fibrils within the f-WT solution incubated for 3 days at 37°C. It also emphasizes that those oligomeric entities might be the membrane-active ones, as they tend to precede the mature fibrils of > 5 nm thickness. To validate this hypothesis, we performed the same real-time experiments, on SLB of diverse controlled composition, after injecting a solution of f-F3A peptides preincubated 3 days at 37°C. Unexpectedly, this mutant, unable to intrinsically form amyloid fibrils (Fig. 1), behaved very similarly as f-WT peptides: it was able to accumulate and elongate as fibrils on eukaryotic mimetic membranes (DOPC/SM/Chol), and more largely on DOPC-containing membranes tested in this study, whereas no deposition or fibrillation was observed on bacterial mimetic membranes (DOPE/DOPG) (Fig. 6A, Fig. S8). Besides, like f-WT, f-F3A preferentially interacts with - and self-aggregates in - the fluid phases of the SLB, as shown by the clustering of elongating fibrils only in DOPC, excluded from the DPPC domains (Fig. 6B, Fig. S8). The transition from small aggregates to thin protofibrils and mature fibrils was finally observed over time, with the former propagating as a soft front potentially inserted within the membrane, and eventually locally disrupting the membrane integrity (Fig. 6B). These nanoscale observations highlight the role of SLB, containing DOPC, as a potential inducer of the aggregation of f-F3A that is otherwise unable to self-assemble in a lipid-free environment. Thus, this mutant cannot serve as a control for the role of intermediate amyloidogenic entities in the potential damage induced by f-WT PSMα3 on SLB.

**Figure 6.**
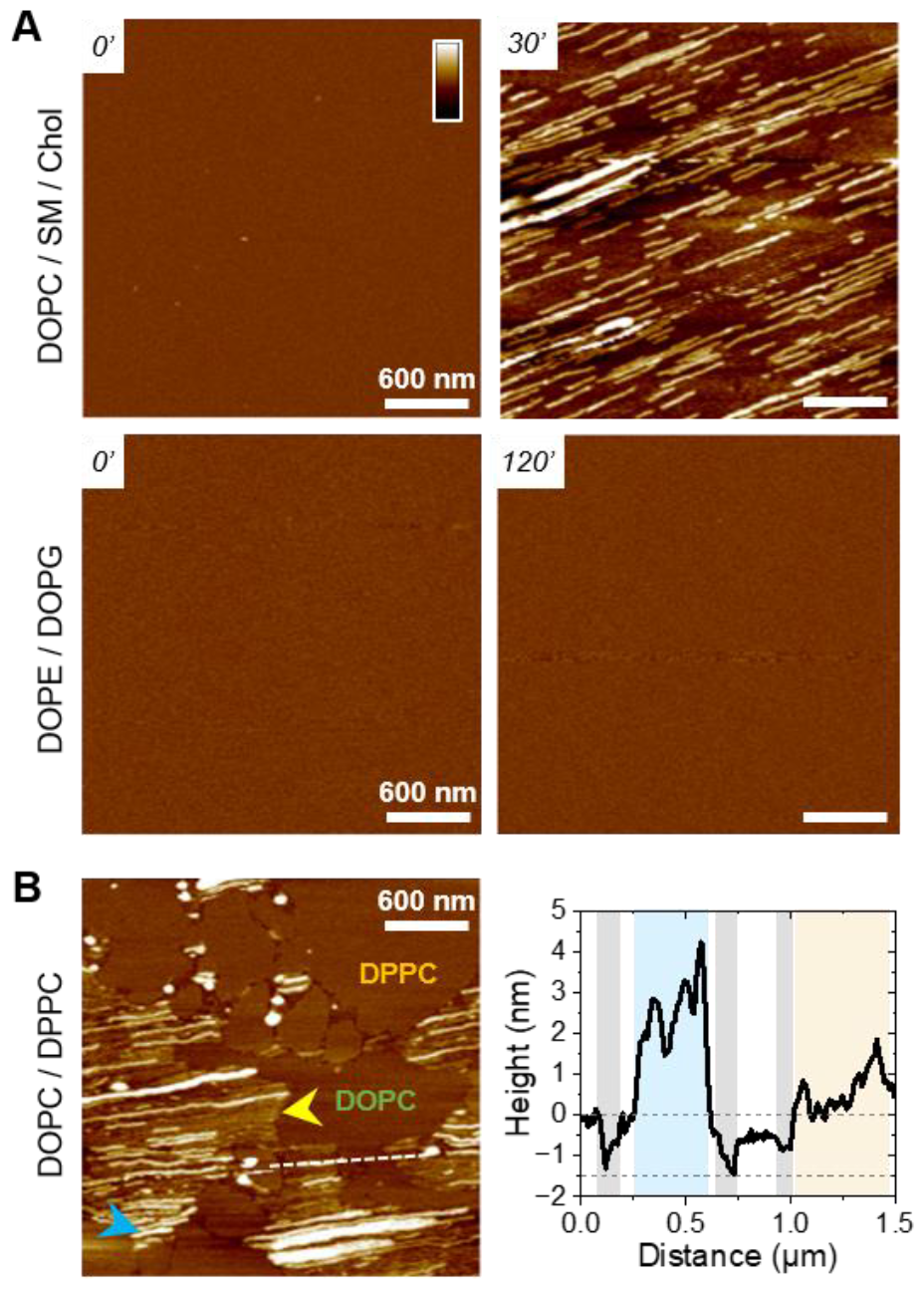
Fibrillation of f-F3A PSMα3 on DOPC-containing membranes. **(A)** AFM topography images of eukaryotic (DOPC/SM/Chol) and bacterial (DOPE/DOPG) mimetic membranes before and after interacting with 5 µM f-F3A. (B) Topographical analysis of a biphasic DOPC/DPPC SLB on which f-F3A has aggregated as thin protofibrils (yellow arrow) and mature fibrils (blue arrow). The height profile along the dashed line notably points to holes in the SLB (grey areas) surrounding the presence of (proto-)fibrils of f-F3A. Color scale bar: 10 nm.

Alternatively, to investigate this point, we carried out experiments where f-WT peptides were injected on the DOPC/DPPC SLB as a monomeric solution (Fig. 7). Instead of pre-incubating the peptides for 3 days at 37°C to favour amyloid fibrillation, and work with this final solution, the f-WT solution has been frozen once prepared to start the experiment only with monomeric entities (f-WT_m). Although the timescale cannot be quantitatively compared, as mentioned above it can differ substantially between different samples, the 3 steps process observed after injecting f-WT on the DOPC-membranes also occurring upon addition of f-WT_m, with similar durations: small aggregates (< 2 nm) first appeared, from which fibrils (∼ 5 nm) start to elongate in time (Fig. 7A-B). Accumulation of f-WT_m was not observed on DOPE/DOPG membranes (data not shown). The aggregates either form small domains, softer than the surrounding membrane and stiffer than the thick fibrils (Fig. 7B), or assemble into thin protofibrils eventually inserted into the membrane, as revealed by holes around them (Fig. 7C). The accumulation of f-WT_m at the interface of DOPC-membranes is further supported by the presence of the Amide I band in ATR-FTIR spectra acquired after a 3 h incubation of f-WT_m on SLB (Fig. 7D). Such infrared experiments also confirm similar impact of the f-WT solutions at 10 µM, either after fibrillation or frozen at the initial monomeric state, on the integrity of SLBs with various controlled composition (Fig. 7E). However, increasing the concentration to 50 µM lead to substantial differences between fibrillated and initial monomeric solution of f-WT (Fig. S9). While both solutions significantly disrupt all membranes, whatever their lipid composition, the deleterious activity of f-WT_m is much reduced on DOPC and DOPE/DOPG SLB compared to f-WT. When SLB are enriched in lipids favouring the formation of gel-like domains (either DPPC or cholesterol), f-WT and f-WT_m similarly and drastically destroy the SLBs. This tend to highlight that (i) entities present over the first 1h (here, the time of SLB treatment by f-WT_m) probably also exist in the final f-WT fibrillated solution (after 3 days at 37°C) and (ii) these entities likely mediate the membrane-deleterious activity of f-PSMα3. Alternatively, one could argue that distinct entities present in f-WT and f-WT_m (*e*.*g*. oligomers *vs*. fibrils) could lead to similar damage on SLB but the real-time AFM observations are not in favour of such hypothesis, as similar morphological evolution of peptides aggregates and protofibrils are observed for both f-WT and f-WT_m. Overall, our AFM and ATR-FTIR results point to a predominant role of f-WT oligomeric, if not monomeric, entities in interacting with supported bilayers, and their preferential aggregation into the liquid phase of these membranes, with the gel-like domains, if present, remaining intact.

**Figure 7.**
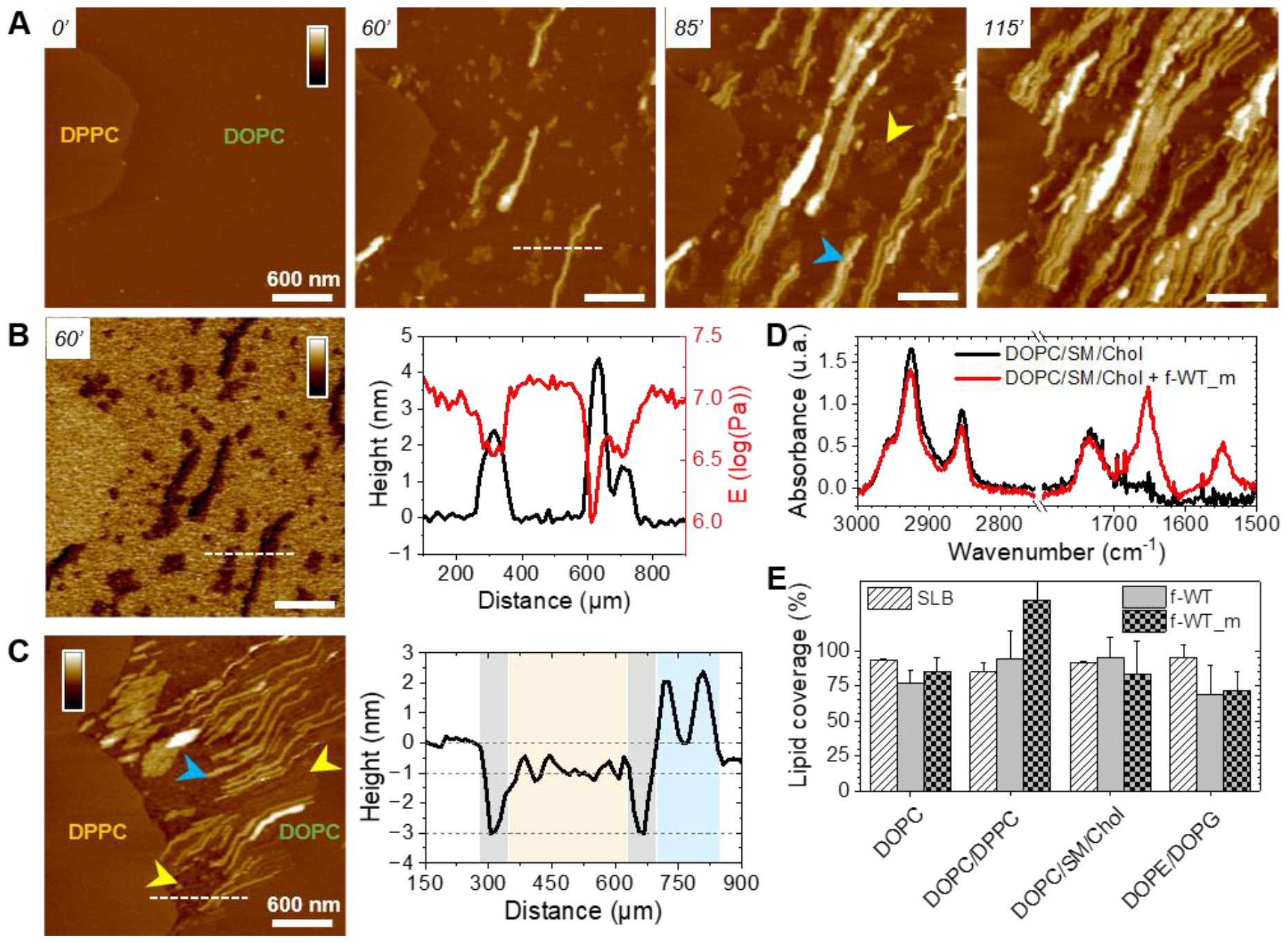
f-WT oligomers are the membrane-active entities. **(A)** AFM topography images, at representative timepoints, of a DOPC/DPPC SLB interacting with a 5 µM monomeric solution of f-WT (f-WT_m). f-WT_m first accumulate as small aggregates (yellow arrows) and fibrillate (blue arrows) only in the fluid DOPC phase. **(B)** DMT modulus image of f-WT_m at the DOPC/DPPC interface, after 60 min of incubation. The height and modulus profiles, along the dashed line, are presented that correlate the presence of a small aggregate and a fibril with the softest areas. **(C)** The thin protofibrils and fibrils eventually co-localize with thinner areas of the membrane (grey areas in the profile). Color scale bars: 20 nm in (A,C), 1,7 log(Pa) in (B). **(D)** ATR-FTIR spectra (p-pol)of a DOPC/SM/Chol SLB before and after a 3 h incubation with f-WT_m at 10 µM, focused on the CH_2_ lipid and Amide bands. **(E)** Comparison of the lipid coverage, obtained in p-pol, of the ATR-FTIR sensor for SLB of diverse composition following a 3 h incubation with 10 µM fibrillated (f-WT) and monomeric (f-WT_m) f-WT solutions. Results are presented as mean ± standard deviation of at least three independent replicates

## Discussion

Combining spectroscopic and high-resolution imaging approaches *in vitro*, we have shown, that both f-WT and f-F3A behave very similarly at the interface of DOPC-containing membranes and bacterial (DOPE/DOPG) membranes. For the former membranes, at low concentration (5-10 µM), both peptides first tend to deposit and accumulate on the membrane as revealed by ATR-FTIR spectroscopy (Fig. 2) and visually confirmed by AFM imaging (Fig. 3, 6). Subsequently, fibrils were observed to grow in time until they eventually form large “carpets” of thin and long fibrils, only in the fluid phase of the membrane. Concomitantly, membrane disruption was locally probed as a thinning effect co-localizing with the growing peptides (Fig. 4, 5). Those local perturbations are consistent with the minor impact of f-WT and f-F3A on DOPC-containing SLB, in terms of lipid depletion (Fig. 2). Significantly, those deleterious effects on membrane integrity are favoured at higher concentration of f-WT (25-50 µM) as illustrated by a decrease in lipid coverage (Fig. S4) and progressive disappearance of the membrane (Fig. S7) following peptide treatment.

### Role of N-formylation in dictating PSMα3-membrane interactions

We have shown that the deposition and fibrillation propensities of formylated (f-) PSMα3 on SLB substantially differ from those of deformylated (df-) PSMα3 previously reported^25^. On one hand, while df-WT forms elongated fibrils at the interface of both eukaryotic (DOPC/SM/Chol) and bacterial (DOPE/DOPG) membranes, f-WT only accumulates and self-aggregates on the former ones. Noteworthy, f-WT and df-WT exhibit similar morphology when elongating on membranes, with a thickness and width of 5 to 10 nm for the thinnest fibrils. On the other hand, while df-F3A forms small aggregates only on DOPE/DOPG SLB, f-F3A displays the exact same behaviour as f-WT with a DOPC-induced fibrillation, and an absence of deposition on DOPE/DOPG SLB. Such differences underscore the role of N-formylation in the mechanism of action of PSMα3 at the cell membrane interface and could arise, in a simplistic view, from the different charges carried by the peptides at the N-terminal when formylated (-C_α_ -NH-COH) or not (-C_α_-NH_3^+^_). Indeed, while electrostatic interactions are favoured between deformylated peptides and negatively charged DOPE/DOPG membranes, they are reduced with the formylated peptides, which might explain the absence of both f-WT and f-F3A accumulation on bacterial membranes (AFM imaging, Fig. 3, 6). Similarly, this reduced charge at the N-terminal would decrease the repulsions between formylated peptides and phosphatidylcholine (PC) groups. While this reasoning can explain the interactions between f-F3A, unlike df-F3A, and DOPC-containing membranes, it cannot account for the similar behaviour of df-WT and f-WT as they both fibrillate on DOPC. This behaviour points to an additional critical role of the phenylalanine in position 3, previously reported in light of PSMα3 toxicity^5,34^, as this aromatic residue would favour the interactions with PC, unlike the alanine present in the F3A mutant. Overall, the different behaviour of f- and df-PSMα3, and WT and F3A, at membrane interfaces, demonstrate that surface induced aggregation of PSMα3 is driven by hydrophobic as well as electrostatic interactions with the lipid acyl chains and their head group, respectively. This lipid-dependent fibrillation is actually shared with pathological amyloids, with kinetic effects (either catalysing or slowing down the aggregation) that relate to the net charge of both lipids and proteins^52^. Such surface-triggered aggregation is also in line with the versatile adsorption and amyloid formation on both hydrophilic and hydrophobic surfaces of PSMα1-4^53^, as well as other bacterial functional amyloids, such as curli proteins CsgA and CsgB of *E. coli*^54^.

### Mechanism of action of PSMα3 at the membrane interface

The “carpet” behaviour that we observe with both f-WT and f-F3A at specific SLB interfaces is usually preceded by local membrane disruption, the extent of which depends on peptide concentration, as above-mentioned, and lipid composition. Indeed, experiments on the binary DOPC/DPPC SLB, that exhibits a clear phase separation between liquid-like DOPC domains and gel-like DPPC domains, have allowed to reveal that both PSMα3 fibrillation and membrane perturbation occur only in the liquid phase of the SLB. Increasing peptide concentration ultimately leads to disruption of both domains. Such phenomena further evidence the role of both electrostatic and hydrophobic interactions for PSMα3 to exert its potential membrane damage. While the ordered state of DPPC acyl chains would prevent insertion of PSMα3, the mismatch between fluid DOPC and gel-like DPPC domains would enhance the hydrophobic interactions with the peptides, thus favouring the nucleation of fibril insertion and elongation within the fluid phase of the membrane from the DPPC edges. Such fluidity-dependent fibrillation and toxicity of amyloids have been previously reported for pathological amyloids. For instance, while fluid POPC or liquid-expanded DPPC (at T > T_m_) facilitates fibrillation of Aβ by favouring a high mobility of the peptides, the liquid-condensed phase of DPPC can retard their self-aggregation^55^, fibrils can even be excluded from such domains in lipid monolayers^56^. Besides, in these fluid membranes, our results point to a membrane perturbation, either as thinning or formation of holes, induced by PSMα3 once a threshold concentration of peptides is reached. This is highly reminiscent of the carpet-like model by which other amphipathic α-helical peptides, the antimicrobial peptides (AMP), permeate cellular membranes^57^ : they first bind to the membrane, cover it and, above a critical density, conformational rearrangement of the peptides within the membrane leads to micelle formation, until the eventual total membrane disruption. Our AFM data further indicate that peptides, notably the thin protofibrils, propagate as a “front” and might be embedded in the membrane, probably in the upper leaflet from the topographical protrusions observed. These observations are partially in agreement with the insertion of df-WT PSMα3 fibrils, but not df-F3A, within the hydrophobic core of DOPC/SM/Chol bilayers^25^. This might also suggest a mode of action for f-PSMα3 that implies transmembrane pore formation, where the peptides could insert perpendicularly in the lipid bilayer. As our ATR-FTIR data do not support a change in membrane fluidity and organization, such pores would result from a barrel-stave pore model, where the amphipathic structure of PSMα3 favours interactions between its hydrophobic residues and the lipid acyl chains and preserve the hydrophilic lipid head group arrangement^58^. Such a mix between a carpet-like and a pore formation models has been described for some AMP that, after accumulating at a critical concentration on bacterial membranes, disrupt their integrity by two-dimensional lateral diffusion within the membrane, illustrated *via* AFM imaging by holes spanning the full or partial (the outer leaflet) bilayer or only a thinning effect^59–61^.

### Role of oligomeric entities in membrane perturbation

Markedly, our AFM data further revealed that membrane thinning, and possible disruption, were usually associated to growing protofibrils that precede the occurrence of mature thick fibrils. Moreover, those phenomena occurred when the peptides (f-WT or f-F3A) were injected either as a monomeric solution (Fig. 7) or after a 3 days-incubation at 37°C to promote amyloid fibrillation (Fig. 4-6). This result first suggests that a reservoir of mono-, more probably oligomeric entities (such as protofibrils) co-exists with the fibrillar entities at the end of the self-assembly process. Besides, these oligomers are likely the membrane-active entities, leading the potential deleterious activities, and eventually the ultimate cytotoxicity. This hypothesis is in agreement with recent evidence, based on mutation assays yielding non-fibrillating peptides or on different levels of cell toxicity among PSMs, that cross-α fibrillation is not sufficient to account for PSMα3 lytic activities^6,22^. Xuan *et al*. also demonstrated, by salt-inducing different PSMα3 assemblies, that oligomers / curvilinear fibrils, and to a lesser extent rigid fibrils, rather than amorphous aggregates, exert high cytotoxicity towards HEK cells *in vivo*^62^. Our *in vitro* observations also highlight that a dynamic exchange between oligomers and fibrils entities, and mutual interactions / aggregation with cell membranes, are required to explain the deleterious activities of both f-WT and f-F3A, in partial agreement with the *in vivo* and *in vitro* behaviour of deformylated PSMα3^22,25,63^. They further revealed, at the nanoscale, that membrane disruption (either thinning or holes that partially span the membrane) is usually locally associated to the presence of oligomers or protofibrils. Interestingly, soluble oligomers of different pathological amyloid proteins were also earlier reported to cause membrane disruption, unlike fibrils, *via* different modes of action: while some exert a detergent-like effect^64^, other induce a membrane thinning due to an increase in conductance without pore formation^65^. Those results thus fuel the debate on the increasingly challenged amyloid cascade hypothesis^40^, according to which insoluble fibrils of pathological amyloids cause cell toxicity^37^.

Despite our above hypothesis and interpretation, it still remains enigmatic why low concentrations of f-WT and f-F3A induced similar effects at the membrane interface, given the reduced toxicity of the deformylated forms of the mutant peptides *in vivo*. To nuance this paradox, one should take into account several points. (1) Despite high cytotoxicity of WT at elevated concentration, not specific to any cell, compared to F3A, both peptides actually exhibit similar lytic activities against HEK cells at concentration lower than 3.5 µM^5^, suggesting that below a critical concentration both peptides could act similarly. (2) Though simplified membrane models are key in understanding fundamental molecular mechanisms *in vitro, in vivo* processes are much more complex with cellular membranes displaying high heterogeneity in the lipid composition, and embedding proteins that could interfere with the lipid pathway. For instance, PSMs are known to bind the formyl peptide receptors 2 (FPR2) on the membrane of immune cells, their complex triggering a host immune cascade^30,32^, and an increased phagocytosis of *S. aureus*^66^. However, PSMs cytotoxicity could be enhanced from the intracellular environment after phagocytosis^23^, thus potentially making interactions of PSMs with such membrane proteins an indirect contribution to the lytic functions of PSMs.

## Conclusions

In conclusion, this work aimed at providing mechanistic and molecular insights into the cytotoxicity of PSMα3, functional amyloids secreted by *S. aureus* pathogens, by investigating their interactions with model biomimetic membranes. Indeed, while the remarkable cross-α fibrillation of PSMα3 has been correlated to its toxicity towards human cells *in vivo*, the molecular mechanisms underlying such function have so far remained elusive, with partial ignorance of the dynamic and mutual interactions at play between PSMα3 self-assembling entities and cell membranes. Here, bridging the behaviour of PSMα3 at the whole membrane level to its local impact on the lipid membrane organization, in real-time and at the nanoscale, we have demonstrated *in vitro* the key roles of N-terminal formylation and intermediate oligomeric entities in leading to membrane damages, likely reflecting PSMα3 lytic activities *in vivo*. Specifically, we have shown that N-formylation, the physiological capping of PSMα3 when initially secreted by *S. aureus*, can modify the peptide binding to cell membranes, in a lipid-dependent manner, by modulating electrostatic interactions with the lipid head charges. In addition, we have revealed that zwitterionic lipids promote the fibrillation of PSMα3 at the membrane interface, only in fluid phases of the bilayer - thus excluded from the ordered and compact domains – as hydrophobic interactions with the acyl chains dictate the potential accumulation and insertion of the peptides in the lipid bilayer. Finally, we have evidenced that oligomeric and protofibrillar structures, rather than mature fibrils, are likely responsible for membrane disruption, *via* membrane thinning and eventual pore formation following peptide accumulation in a “carpet” fashion. Such findings, beyond highlighting the critical importance of N-formylation in PSMα3-membrane interactions along with its *in vivo* relevance, thus additionally fuel the increasing debate on the amyloid cascade hypothesis, even in the context of functional amyloids.

## Materials and methods

### Materials

Formylated PSMα3 peptides, in the WT (f-MEFVAKLFKFFKDLLGKFLGNN) and mutant F3A (f-ME<u>A</u>VAKLFKFFKDLLGKFLGNN) forms, were purchased from GenScript at ≥ 98 % purity. Mass spectrometry experiments were performed to confirm the purity of the peptides. DOPC, DPPC, DOPE, DOPG, cholesterol (Chol, ovine wool, > 98 %), and sphingomyelin (SM, brain porcine) were purchased from Avanti Polar Lipids, trifluoroacetic acid (TFA, ≥ 99 % HPLC grade) from Fisher Scientific and thioflavin T (ThT) and hexafluoroisopropanol (HFIP) from Sigma Aldrich.

### Peptide preparation

f-WT and f-F3A solutions were prepared by dissolving the peptide powder at a concentration of 1 mM in a (1:1) mixture of HFIP/TFA, for 1 h at room temperature (RT). Solvent was evaporated under a stream of dry N_2_ and solvent residues were evaporated under vacuum in a dessicator for 2 h. The resulting peptide film was rehydrated with ultrapure water at a concentration of 1 mM, on ice, and sonicated for 5 min. This solution was then further diluted, at the desired concentration, in a 10 mM sodium phosphate buffer complemented with 150 mM NaCl (pH 8.0). It was finally centrifuged (10 000 rpm, 5 min, 4°C), and the supernatant was collected to avoid initial aggregates. This final peptide solution was either fast-frozen in liquid nitrogen, and kept at -80°C for investigation on monomeric peptide solution, or incubated for 3 days at 37°C under gentle agitation (∼400 rpm).

### Thioflavin T (ThT) fluorescence assay

The kinetics of PSMα3 self-aggregation was monitored using the variations of the fluorescence intensity of ThT dye (λ_excitation_ = 449 nm / λ_emission_ = 482 nm). Fluorescence measurements were performed on a CLARIOstar plus (BMG Labtech) plate reader, using standard 96 or 384 well flat-bottom black plates, sealed with a transparent cover sticker to avoid evaporation. Assays were performed 3 times independently, each in triplicate, in a final volume of 200 or 50 µL, for 96 and 384 well plates respectively, containing (i) 200 µM ThT and appropriate volumes of (ii) buffer (10 mM sodium phosphate buffer complemented with 150 mM NaCl (pH 8.0)) and (iii) peptides to reach the desired concentration (from 10 to 100 µM). The peptides were freshly prepared as described above, prior to ThT fluorescence assays, to avoid aggregation preceding the first measurements. The ThT solution was obtained by diluting a stock of ThT at 10 mM in water into the appropriate buffer at a final concentration of 1 mM. The fluorescence intensity was measured at 37°C, with a 500 rpm orbital shaking for 15 s before each cycle, with up to 1000 cycles of 5 min each.

### Transmission electron microscopy (TEM)

TEM was performed on peptide solutions after 3 days of incubation at 37°C (following either ThT fluorescence assays or incubation “on bench” under similar conditions), to assess the formation of aggregates and amyloid fibrils for each peptide solution prepared. 4.2 µL of the peptide solution (50-100 µM) were adsorbed onto glow-discharged carbon coated 300 mesh copper grids for 2 min and the excess solution was blotted with filter paper. Negative staining was then performed using a 1% uranyl acetate solution applied to the grids for 30 s and blotted again for the grids to dry. The grids were finally examined using a CM120 electron microscope operating at 120 kV with an LaB6 filament. Different areas of the grids were images to get a representative picture of the entities (f-WT and f-F3A) formed over the incubation.

### Supported lipid bilayer (SLB) preparation

Phospholipid vesicles were prepared as followed. DOPC, DPPC, DOPG, DOPE, SM, and Chol dissolved in chloroform were pipetted into an Eppendorf in appropriate volumes to reach a molar ratio of (67:8:25) for DOPC/SM/Chol, (1:1) for DOPE/DOPG and (1:1) for DOPC/DPPC and stirred to get homogeneous solutions. The solvent was then removed first by evaporation under a stream of dry N_2_ and then by placing the ependorf under vacuum in a desiccator for 2 h. The lipid films were then rehydrated with the same buffer as the one used for peptide preparations (10 mM sodium phosphate buffer complemented with 150 mM NaCl (pH 8.0)) to form multilamellar vesicles (MLV) at a final concentration of 2 mg/mL. This MLV suspension was then sonicated over 3 cycles of 10 min., with an amplitude of 40 % and 3 s pulses to obtain small unilamellar vesicles (SUV). While sonicating, the ependorf was kept in an ice bath to limit heating. The SUV suspension was finally filtered on 0.2 µm filters to remove eventual residues from the ultrasound probe, and stored at 4°C, before any experiment, for maximum 2 weeks.

From those SUV suspensions, SLB were formed on the appropriate substrate according to the vesicle fusion method. For AFM experiments, 100 µL of a 1 mg/mL SUV suspension were applied for 30 min. on a freshly cleaved mica disk, heated to 60°C if needed (*e*.*g*. for the binary mixture DOPC/DPPC). Attention was paid to avoid dewetting: buffer was added regularly for the SLB to always stay hydrated. In case of heating, the sample was slowly cooled down to RT for 30 min. Whatever the temperature of incubation, the samples were finally heavily and carefully rinsed with the buffer (∼ 10 times, V = 100µL) to remove all unabsorbed vesicles. 80µL of buffer were finally added onto the SLB for AFM investigation in liquid. For ATR-FTIR experiments, 20 µL of a 1 mg/mL SUV suspension were applied directly on an ATR germanium crystal hosted in a homemade liquid chamber. After 5 min., the SLB was rinsed 6 times with buffer, letting a final volume of 20 µL for infrared spectra acquisition. For both AFM and ATR-FTIR experiments, when working with anionic lipids (*e*.*g*. DOPG), 1-2 mM of CaCl_2_ was added to the buffer to favour SUV fusion on the substrates.

### Attenuated total reflectance Fourier Transform infrared spectroscopy (ATR-FTIR)

ATR-FTIR spectra were recorded, in buffer conditions (V = 20 µL) at room temperature (20 °C), on a Nicolet iS50 FTIR spectrometer equipped with an MCT detector cooled with liquid N_2_, and an ATR accessory mounted with a germanium crystal (one reflection). Polarized spectra (incident light at 0° and 90°, *i*.*e. s-* and *p-*polarizations respectively) were recorded with 200 scans and a spectral resolution of 2 cm^-1^. To remove the contribution of ambient air and buffer, background and buffer spectra were first collected for both *s-* and *p-*polarizations, and then subtracted from all samples spectra either during acquisition or in post-processing. SLB was then formed on the germanium crystal, and rinsed as described above. Spectra in *s-* and *p-*polarizations were collected before peptide addition, or after 3 h to assess its stability over time. Provided the correct formation and stability of the SLB, the peptide of interest was then injected in the final volume of 20 µL at a final concentration of 50 or 10 µM for 1 or 3 h respectively. Spectra following peptide-membrane interactions were collected after rinsing the liquid chamber, to avoid the contributions of non-adsorbed peptides and lipids in solution close to the germanium crystal. ATR-FTIR spectra were post-processed using the Omnic software, to substract the buffer contribution and correct the baseline at the following points: 3500, 3000, 2800, 1800, and 1000 cm^-1^. Deconvolution of the Amide I band was performed with OriginPro (OriginLab). Every experiment was performed at least 3 independent times. For clarity, only spectra obtained in the *p-*polarization (and resulting analysis) are presented herein, as the same variations following peptide-membrane interactions were observed in the *s-*polarization and the *p-pol* spectra were the most intense.

### Atomic force microscopy (AFM)

AFM imaging of the SLBs was performed using the PeakForce Quantitative Nano-Mechanics (PF-QNM) mode on an Dimension Fast-Scan setup (Bruker) in buffer conditions at room temperature (20 °C). Nitride-coated silicon cantilevers (SNL-C, Bruker) with a nominal spring constant of 0.24 N/m, and a tip radius of 2 nm, were used and calibrated before any experiment using the thermal noise method. The images, analysed and processed with the Nanoscope Analysis software (Bruker), were acquired with a scan rate of ∼1 Hz, a Peakforce amplitude of 50 nm and a Peakforce frequency of 1 kHz, and the applied force kept as low as possible to minimize any tip-induced damage (< 1 nN). Once the formation and stability of the SLB have been confirmed by AFM imaging, the peptides of interest were injected at a final concentration of 5 µM, in a total volume of 80 µL, if not otherwise stated. The same area (typically 3x 3 µm^2^) was then imaged in real time with an approx. 1 image every 5 min. to probe and correlate eventual peptide aggregation and membrane damage. Zoom out and different areas were also scanned to ensure that those phenomena are not zone-dependent, or due to tip scanning artefacts. Of note, along with the topographic images, AFM forces curves were recorded in each pixel of the scanned area, and mechanical images of the areas were simultaneously recorded. In this study, we present the Derjaguin, Muller, Toropov (DMT) modulus map of the areas of interest. This DMT modulus (E) is obtained by fitting the retract curve with the following model:

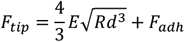

Where F_tip_ is the force on the tip, F_adh_ the adhesion force if existent, R the tip radius, and d the tip-sample distance.

Every experiment was performed at least 3 independent times, and the images presented herein are representative of the obtained results.

## Author contributions

LB, KJ and MMG collected and analysed all experimental data (ThT, TEM, ATR-FTIR and AFM). AV and MMG supervised and trained students in AFM investigation. MMG conceived the project for which she obtained national and European funding. LK, SL, CF, MM and MMG designed the experimental methodology, and interpreted experimental results. All authors contributed to the manuscript writing.

## Conflicts of interest

There are no conflicts to declare

## Acknowledgements

The authors thank Sisareuth Tan (assistant engineer, CNRS) and Sandrine Villette (research engineer, CNRS) for the training and technical help in TEM (IECB platform (UAR3033)) and ATR-FTIR experiments respectively. They thank Katell Bathany (research engineer, University of Bordeaux) for the mass spectrometry experiments that confirmed the quality of the peptides used within this study. They finally thank the VIbrAFM platform on which the AFM and ATR-FTIR experiments were performed. MMG thanks the *Fondation pour la Recherche Médicale* (project ARF202110014167) and the European Union and the Marie Skłodowska-Curie Actions for funding the PSMNano project (grant n° 101064573). Views and opinions expressed are however those of the authors only and do not necessarily reflect those of the EU. Neither the EU nor the granting authority can be held responsible for them.

## Notes

### Competing Interest Statement

The authors have declared no competing interest.

