## Supplementary figures for "N-formylation modifies membrane damage associated to PSMα3 interfacial fibrillation"

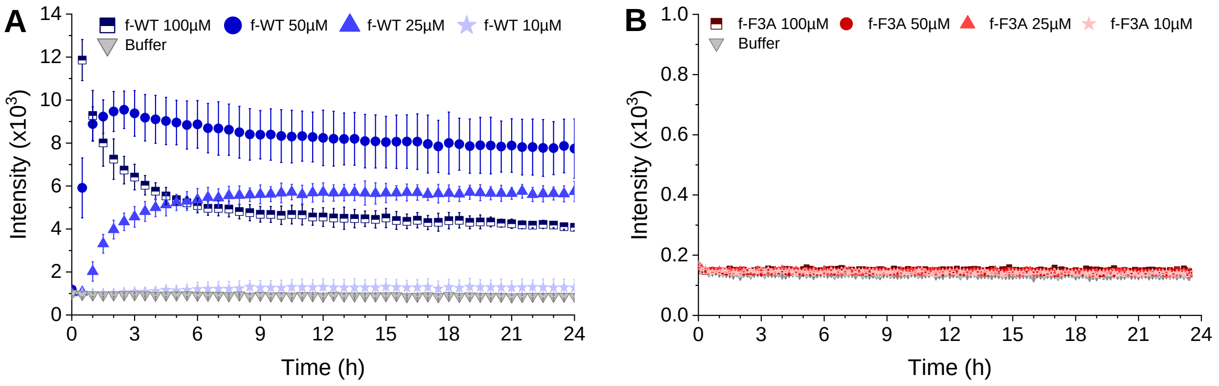


**Figure S1.** Kinetics of **(A)** f-WT and **(B)** f-F3A fibrillation followed by ThT fluorescence, at 37°C and varying peptide concentrations. Error bars stand for the standard deviation between three replicates.

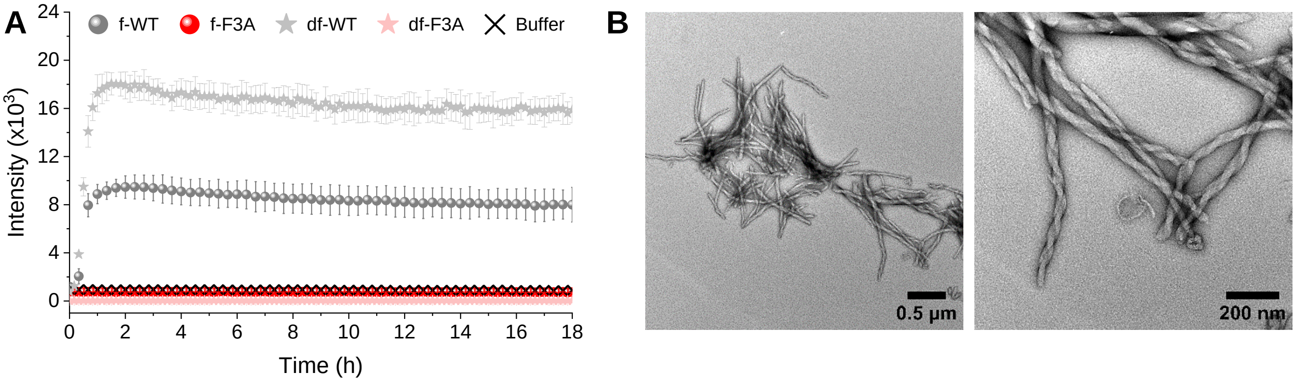


**Figure S2. (A)** Comparison of the fibril formation of formylated (f-) and deformylated (df-) PSMα3 followed by ThT fluorescence (37°C, peptide concentration 50 µM). Error bars stand for the standard deviation between three replicates. **(B)** Negatively stained TEM images of df-WT fibrils obtained after 3 days incubation at 37°C.

**A**

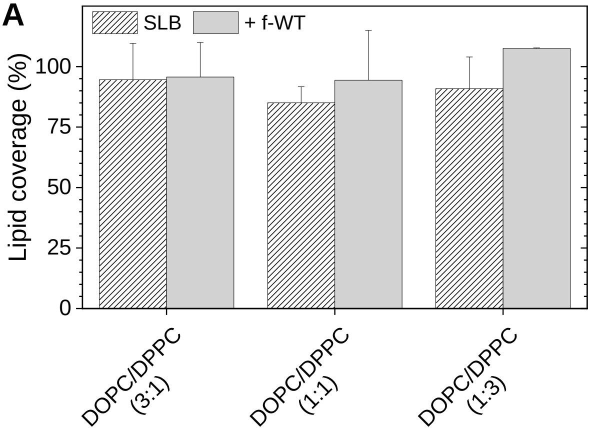


**B**

| | f-WT | $\nu_{as}$ ($CH_2$) ($cm^{-1}$) | $R_{ATR}$ ($\nu_{as}$ ($CH_2$)) |
|---|---|---|---|
| **DOPC / DPPC (3:1)** | - | 2922 | 1.26 ± 0,11 |
| | + | 2923 | 1.26 ± 0.07 |
| **DOPC/DPPC (1:1)** | - | 2920 | 1.20 ± 0.08 |
| | + | 2921 | 1.36 ± 0.15 |
| **DOPC/DPPC (1:3)** | - | 2919 | 1.35 ± 0.13 |
| | + | 2919 | 1.35 ± 0.08 |

**Figure S3. (A)** Lipid coverage of the ATR-FTIR sensor, determined *via* the variations in $\nu_{as}$ ($CH_2$) intensity, for binary DOPC/DPPC SLB, with varying concentration of DPPC, and following f-WT addition (3 h, 10 µM). Results are presented as mean ± standard deviation of at least three independent replicates. **(B)** Table of the wavenumber and dichroic ratio of the $CH_2$ of DOPC/DPPC membranes with varying ratio of DPPC, before (-) and after (+) f-WT addition at 10 µM for 3h. Analyzed spectra were obtained in the *p*- polarization.

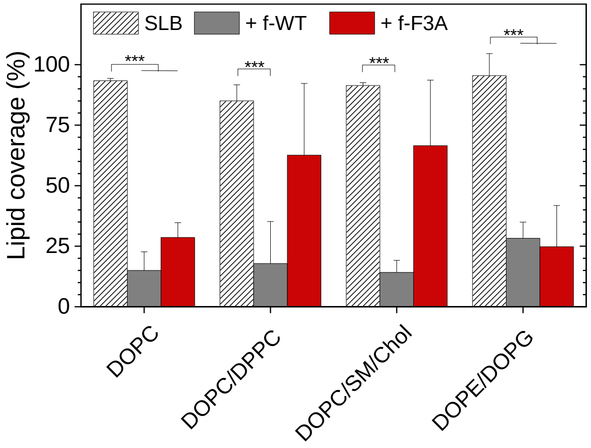


**Figure S4.** Lipid coverage of the ATR-FTIR sensor, determined *via* the variations in $v_{as}$ ($CH_2$) intensity, for SLB of diverse composition following PSMα3 incubation for 1 h at 50 µM. Results are presented as mean ± standard deviation of at least three independent replicates. *** $p < 0.001$. Analyzed spectra were obtained in the *p*-polarization.

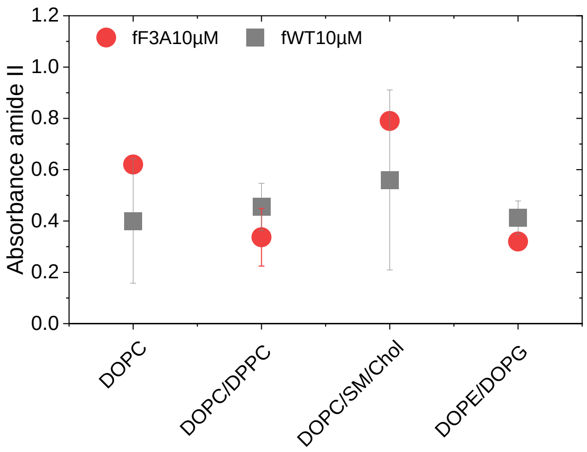


**Figure S5. Quantification of PSMα3 deposition at the SLB interface.** Absolute intensities, in the *p-pol*, of the Amide II band (~ 1550 $cm^{-1}$) after a 3h incubation of PSMα3 (f-WT and f-F3A) on SLB of different compositions.

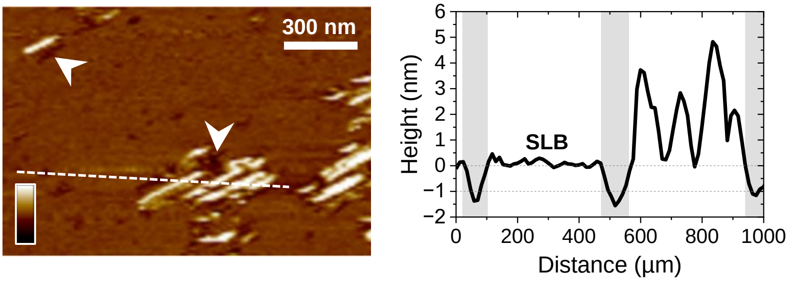


**Figure S6.** Topographical analysis of a DOPC/SM/Chol membrane interacting with f-WT (5 µM). The AFM topography image and the corresponding height profile along the dashed line showcase short fibrils (~ 5 nm) surrounded by areas thinner than the SLB (white arrows and grey area). Small holes can also be observed in the membrane, independently of the f-WT presence. Color scale bar: 10 nm.

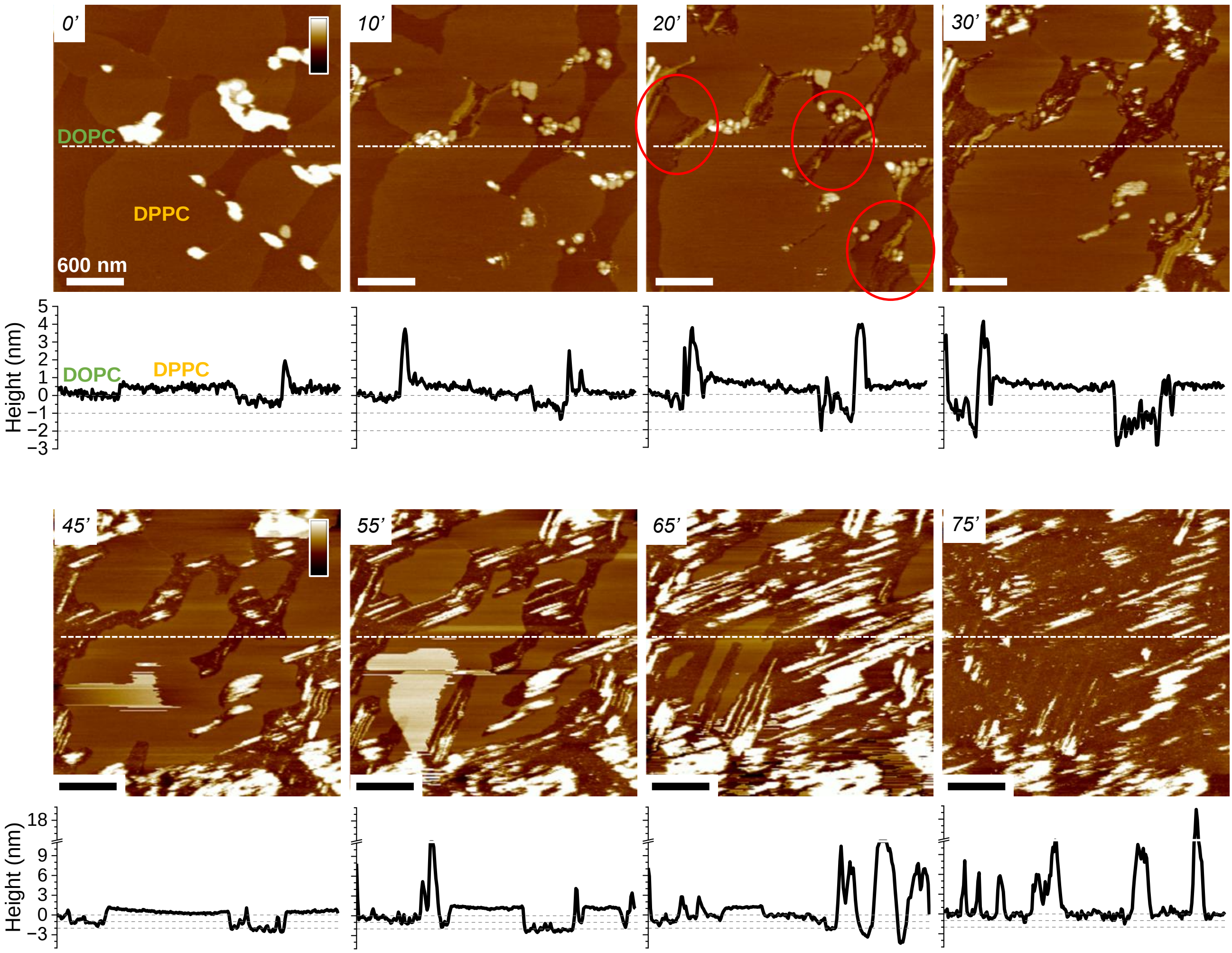


**Figure S7.** Topographical analysis of a DOPC/DPPC membrane interacting with f-WT at a concentration of 15µM, at representative timepoints. Height profiles along the dahsed lines in the AFM topography images are shown to quantitatively measure membrane damage and peptide aggregation. Red circles point to areas where membrane thinning starts occuring (from 1 to 2 nm), as also shown by the height profiles over time.

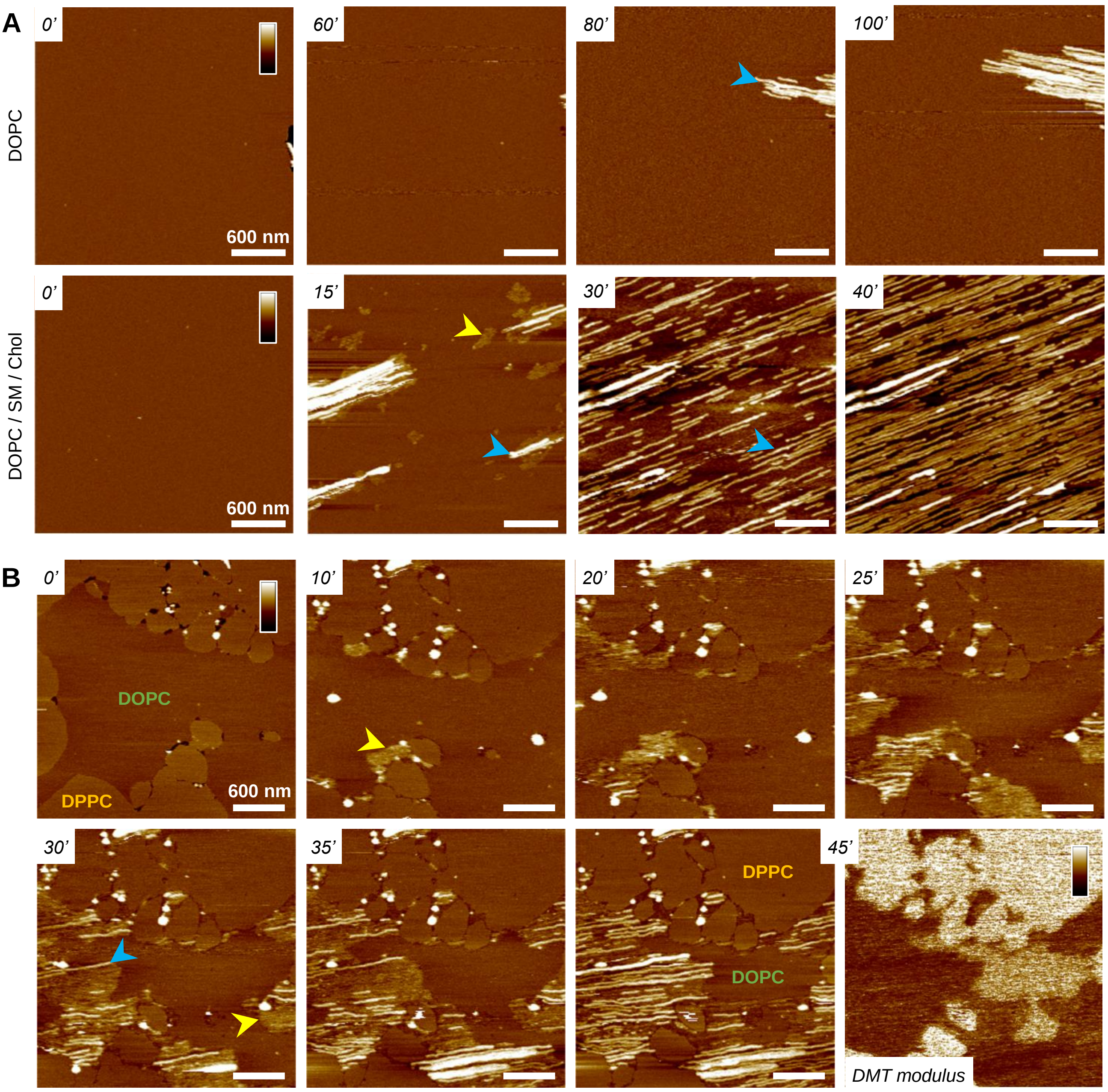


**Figure S8. Fibrillation of f-F3A PSMα3 on DOPC-containing membranes. (A)** AFM topography images, at representative timepoints, of various DOPC-containing SLB interacting with 5 µM f-F3A. Small aggregates (yellow arrows) first appear before elongating as thin (blue arrows) fibrils. **(B)** Timelapse of the growth of both protofibrils (yellow arrows) and mature fibrils (blue arrows) of f-F3A in the fluid DOPC phase of a binary DOPC/DPPC SLB. The front of protofibrils is further evidenced by their mechanical properties (DMT modulus image at 45min of incubation), softer than the surrounding membrane. Color scale bars: 10nm, 2 log(Pa).

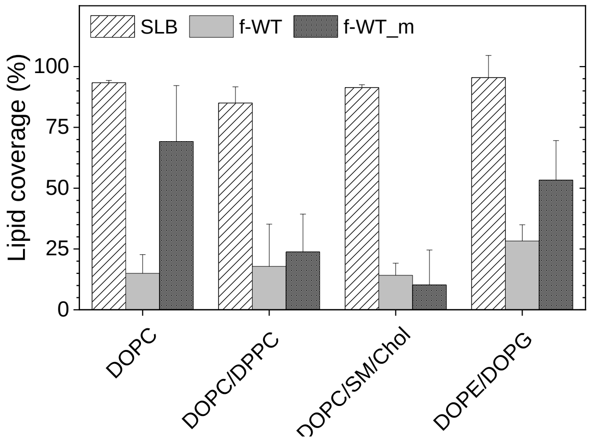


**Figure S9.** Lipid coverage of the ATR-FTIR sensor, determined *via* the variations in $v_{as}$ ($CH_2$) intensity, for SLB of diverse composition following f-WT incubation at 50 µM for 1 h, injected as a fibrillated solution (3 days at 37°C, f-WT) or an initial monomeric solution (f-WT_m). Results are presented as mean ± standard deviation of at least three independent replicates. Analyzed spectra were obtained in the *p-* polarization.
